# Low light intensity delays vegetative phase change in *Arabidopsis thaliana*

**DOI:** 10.1101/2021.01.08.425922

**Authors:** Mingli Xu, Tieqiang Hu, R. Scott Poethig

## Abstract

Plants that develop under low intensity light (LL) often display a phenotype known as the “shade tolerance syndrome (STS)”. This syndrome is similar to the phenotype of plants in the juvenile phase of shoot development, but the basis for this similarity is unknown. We tested the hypothesis that the STS is regulated by the same mechanism that regulates the juvenile vegetative phase by examining the effect of LL on rosette development in *Arabidopsis thaliana*. We found that LL prolonged the juvenile vegetative phase and that this was associated with an increase the expression of the master regulators of vegetative phase change, miR156 and miR157, and a decrease in the expression of their SPL targets. Exogenous sucrose partially corrected the effect of LL on seedling development and miR156 expression. Our results suggest that the response of Arabidopsis to LL is mediated by an increase in miR156/miR157 expression and by factors that repress SPL gene expression independently of miR156/miR157, and is caused in part by a decrease in carbohydrate production. The effect of LL on vegetative phase change does not require the photoreceptors and transcription factors responsible for the shade avoidance syndrome, implying that light intensity and light quality regulate rosette development by different pathways.

## Introduction

Over the last 50 years, research on the response of plants to light has largely focused on the effect of the presence vs. absence of light and the effect of light quality on early seedling development (Smith, 1982; Chory, 1997; Roig-Villanova and Martinez-Garcia, 2016; Ballare and Pierik, 2017). Extensive research on the effect of red, far red, and blue light on hypocotyl elongation in Arabidopsis has provided a deep understanding of the molecular mechanism by which specific wavelengths of light regulate this developmental process. In contrast, the effect of light intensity on plant development has received relatively little attention, despite the fact that in nature plants can experience dramatic variation in light intensity on both short (e.g. seconds or minutes) and long (hours, months, years) time scales.

Plants adapted for growth under low light intensity (LL), as well as shoots exposed to a prolonged period of LL, often exhibit a phenotype known as the shade tolerance syndrome (STS) (Valladares and Niinemets, 2008). This phenotype consists of a large number of physiological, morphological, and anatomical traits that optimize carbon gain; indeed, it is generally assumed that the primary, if not the only, function of the STS is to maximize photosynthesis under LL. However, some of the anatomical and morphological traits associated with the STS have no known function in photosynthesis (Valladares and Niinemets, 2008), raising the question of why the expression of these traits is associated with LL. Additionally, many of the anatomical/morphological features associated with the STS are distinct from, and in some cases opposite to, the traits that characterize the shade avoidance syndrome (SAS) (Valladares and Niinemets, 2008; Ballare and Pierik, 2017), a phenotype generated by a low R:FR light ratio or exposure to FR light near the end of the day. This is significant because in nature LL is typically caused by the presence of surrounding plants, which both decrease light intensity and decrease the R:FR ratio by absorbing red light during photosynthesis (Chory, 1997). This suggests that plants respond to LL and a decreased R:FR ratio by different mechanisms, and are able to “choose” the mechanism that is appropriate for the specific light conditions in which they find themselves.

It has been recognized for over a hundred years that leaves grown in LL resemble leaves produced during the juvenile phase of shoot development (Nordhausen, 1912; Schramm, 1912; Njoku, 1956). However, the basis for this similarity is unknown. One possibility is that the LL delays the juvenile-to-adult transition (vegetative phase change) or reactivates the juvenile vegetative phase in adult plants. An alternative possibility is that low light intensity affects some of the traits associated with the juvenile phase, without actually affecting the timing of vegetative phase change (Jones, 1995, 1999). Both hypotheses imply that the STS is regulated in whole or in part by the vegetative phase change pathway, and recent evidence that juvenile leaves are photosynthetically more efficient under low light intensity than adult leaves (Lawrence et al., 2020) provides intriguing support for this hypothesis. To investigate the role of the vegetative phase change pathway in the STS, we characterized the effect of LL on vegetative phase change in Arabidopsis.

Vegetative phase change is regulated by a decline in the level of the related miRNAs, miR156/157, and the resulting increase in the expression of their targets, SQUAMOSA PROMOTER BINDING PROTEIN-LIKE (SPL) transcription factors (Wu and Poethig, 2006; Chuck et al., 2007; Wu et al., 2009). In Arabidopsis, miR156/157 directly regulate 10 *SPL* genes (Rhoades et al., 2002; Xu et al., 2016; He et al., 2018). These *SPL* genes are responsible many aspects of adult shoot development, including adult leaf morphology, abaxial trichome production, reduced branching, a low propensity for adventitious root initiation and shoot regeneration, low anthocyanin production, decreased insect resistance, and increased thermosensitivity (Chuck et al., 2007; Schwarz et al., 2008; Wang et al., 2009; Wu et al., 2009; Yamaguchi et al., 2009; Yu et al., 2010; Gou et al., 2011; Padmanabhan et al., 2013; Stief et al., 2014; Zhang et al., 2015; Xu et al., 2016; Mao et al., 2017; Nguyen et al., 2017).

Approximately 90% of the miR156 in the rosette leaves of Arabidopsis is produced by *MIR156A* and *MIR156C* (He et al., 2018). The down-regulation of these genes in successive rosette leaves is largely dependent on changes in their chromatin structure (Pico et al., 2015; Xu et al., 2016; Xu et al., 2018). Early in shoot development, *MIR156A*/*MIR156C* contain elevated levels of the active histone modification, H3K27ac (Xu et al., 2016), as well as high levels of H2A.Z, which facilitates the deposition of the active histone modification, H3K4me3 (Xu et al., 2018). The subsequent repression of these genes is mediated by the deposition of H3K27me3, in a process modulated by PRC1 (Pico et al., 2015), PRC2 (Xu et al., 2016), and the chromatin remodelers PICKLE (Xu et al., 2016) and BRAHMA (Xu et al., 2016). In addition to being post-transcriptionally repressed by miR156/157, the expression of *SPL* genes is regulated by several processes that operate independently of miR156/157, including H2A monoubiquitination (H2Aub) by AtRING1A/AtRING1B (Kim et al., 2015; Li et al., 2017), histone acetylation by SAGA-like histone acetyletransferase (HAG1) (Kim et al., 2015), and O-linked N-acetyleglucosamine modification of SPL proteins(Xu et al., 2017). However, it is currently unknown if these modifications control the temporal and spatial pattern of *SPL* gene expression.

Most of the research on the response of Arabidopsis to shade has focused on the mechanism of the SAS (Casal, 2013; Roig-Villanova and Martinez-Garcia, 2016; Ballare and Pierik, 2017). As noted above, this syndrome is produced by exposure to FR-enriched light and a decrease in the intensity of blue light, leading to an increase in the expression of the transcription factors PIF4, 5 and 7, the primary mediators of this syndrome. It has been suggested that some aspects of the SAS are mediated by the miR156/SPL pathway based on the observation that miR156 levels decline by 80% while SPL transcript levels increase up to 4-fold when plants are briefly exposed to FR light at the end of the day (Xie et al., 2017). However this result has yet to be reconciled with the observation that the primary sources of miR156 in Arabidopsis—*MIR156A* and *MIR156C*—are not targets of the PIF transcription factors that mediate this EOD FR response (Xie et al., 2017). Here, we show that the juvenile-to-adult transition in Arabidopsis is delayed by LL, and that this response is mediated by both an increase in the level of miR156/miR157, and by a miR156-independent decrease in the level of SPL transcription factors. Phytochrome- and cryptochrome-mediated signaling are not required for this response, suggesting that the response of plants to light quality and their response to LL are mediated by different regulatory pathways. Along with the evidence that juvenile leaves are photosynthetically more efficient under LL than adult leaves (Lawrence et al., 2020), these results suggest that the STS is under the regulation of the miR156/SPL vegetative phase change pathway.

## Results

### Vegetative phase change is delayed by LL

To study the role of light intensity in vegetative phase change without the complicating effect of a difference in flowering time, we grew wild-type Col in short days under HL (180 µmol m^-2^ s^-1^) and LL (62 µmol m^-2^ s^- 1^) conditions. This was accomplished by varying the number of fluorescent light bulbs in otherwise identical growth chambers. Plants grown in LL were lighter green and slower to develop than plants grown in HL (Fig. 1A, B), and their leaves were rounder and less serrated than HL plants (Fig. 1C). Additionally, abaxial trichome production was significantly delayed in LL plants compared to plants grown in HL (Fig. 1C). These results demonstrate that LL delays vegetative phase change.

**Figure 1.**
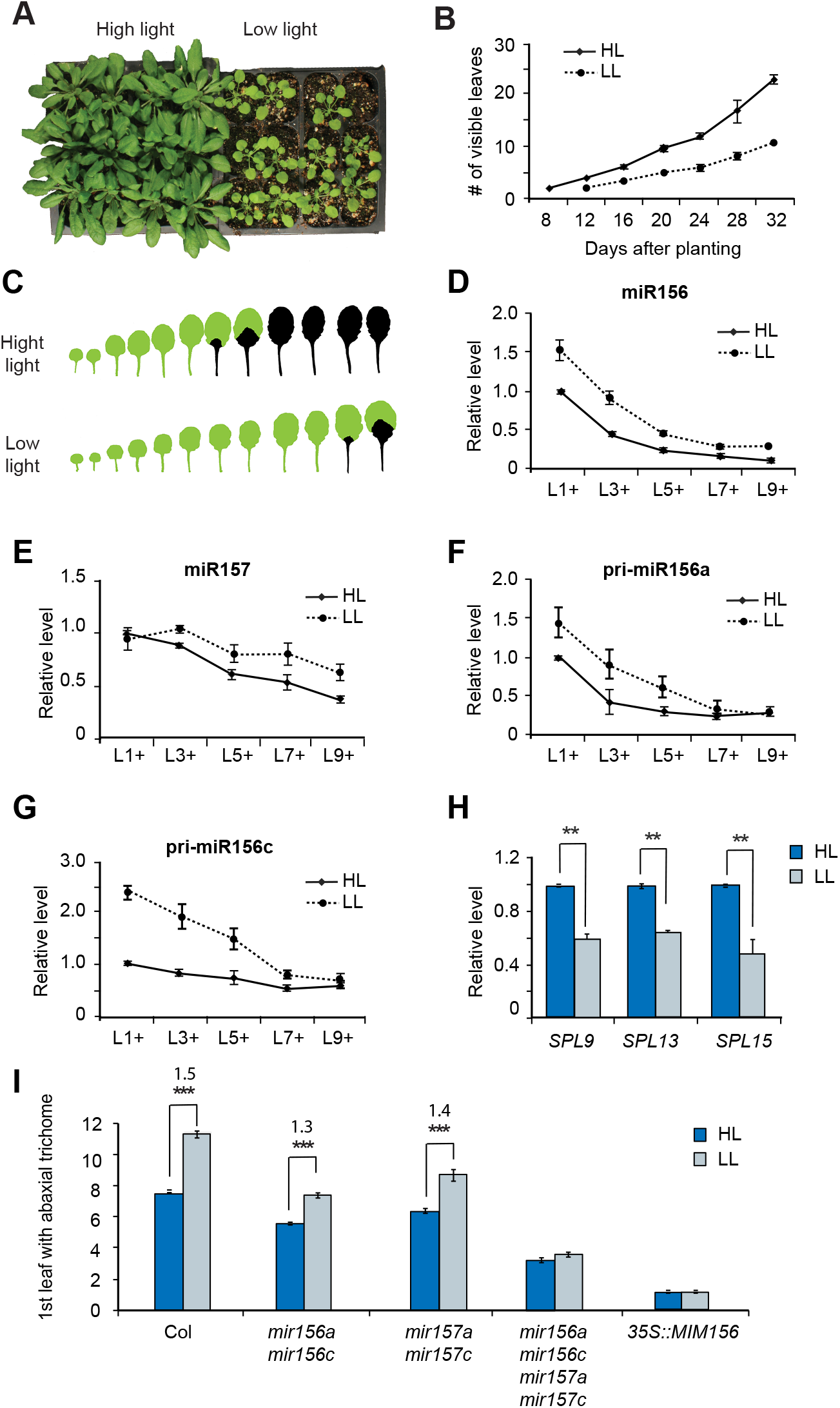
LL delays vegetative phase change via miR156/157. (A) Col plants grown under HL (left) are greener and have more leaves than plants grown under LL (right). (B) Rate of leaf initiation in Col grown in HL and LL. n = 32 for each condition. P<0.001, one-way ANOVA. (C) LL delays the production of rosette leaves with abaxial trichomes in Col. Green = no abaxial trichomes; Black = abaxial trichomes. (D-G) RT-qPCR of miR156 (D), miR157 (E), pri-miR156a (F), and pri-miR156c (G) in the shoot apices of Col plants grown in HL and LL. Samples were harvested according to their developmental stage. (H) RT-qPCR analysis of the transcripts of miR156/157-targeted genes in L3+ shoot apices. In D-H, values are the mean ± SEM from 3 biological replicates. (I) The first leaf with abaxial trichomes in Col and mutant plants deficient for miR156, miR157, and miR156 & miR157 under HL and LL conditions. n = 20 to 24 for each genotype. ** denotes p<0.01, and *** denotes P<0.001, one-way ANOVA.

To determine if the effect of light intensity on vegetative phase change is attributable to changes in the level of miR156/157, we measured the abundance of these miRNAs in plants grown in short days (SD) under HL and LL. Because plants grow more slowly in LL (Fig. 1B), we sampled shoot apices according to their developmental stage. Samples were collected when leaves 1, 3, 5, 7, or 9 were 1mm in length, and consisted of this leaf primordium and all younger leaf primordia and the shoot apical meristem (L1+, L3+, L5+, L7+ and L9+).

The temporal pattern of miR156 expression was similar in HL and LL, but the amount of miR156 in LL shoot apices was significantly higher than the amount of miR156 in HL apices at every stage of development (Fig. 1D). There was no significant difference in the abundance of miR157 in L1+ apices grown in LL and HL. In HL, the abundance of miR157 began to decline at L3+, whereas in LL it began to decline at L5+ (Fig. 1E). This slight delay in the down-regulation of miR157 relative to miR156 suggests that light intensity regulates the expression of miR156 and miR157 by different mechanisms. We then analyzed the effect of LL on the primary transcripts of *MIR156A* and *MIR156C*, the major sources of miR156 in Arabidopsis (He et al., 2018). Both transcripts were present at significantly higher levels at early stages of development (L1+, L3+, L5+) in LL-than in HL-grown plants (Fig. 1F and G), but declined to the same level in older (L9+) plants. This result is consistent with the expression pattern of the mature miR156 transcript, and suggests that these genes are primarily responsible for this expression pattern. Consistent with the elevated levels of miR156, the SPL9, SPL13, and SPL15 transcripts were less abundant in LL than in HL plants (Fig.1H and Fig. S1A).

To determine if miR156 and miR157 are responsible for the delay in vegetative phase change under LL, we compared the phenotype of stocks deficient for miR156 (*mir156a mir156c*) or miR157 (*mir157a mir157c*), and stocks deficient for both of these miRNAs (*mir156a mir156c mir157a mir157c* and *35S::MIM156*) under HL and LL. Abaxial trichome production was significantly delayed by LL in both wild-type Col and in the *mir156a mir156c* and *mir157a mir157c* double mutants. However, LL had a slightly smaller effect on abaxial trichome production in these mutants than in Col, and had no significant effect on abaxial trichome production in the *mir156a mir156c mir157a mir157c* quadruple mutant or in *35S::MIM156* (Fig. 1I). These results demonstrate that miR156 and miR157 both contribute to the effect of LL on abaxial trichome production.

### A miR156-independent pathway is involved in the LL response

Although LL did not affect abaxial trichome production in the *mir156a mir156c mir157a mir157c* and *35S::MIM156* lines (Fig. 1I), it did have a significant effect on leaf morphology in these miR156/157 deficient lines (Fig. 2A; Fig. S1B). In Arabidopsis, juvenile leaves have smaller angle between the petiole and the leaf blade than adult leaves (Yang et al., 2011; He et al., 2018). We found that LL significantly decreased the leaf blade: petiole angle of leaves 1&2 in both Col and in mutants deficient for miR156/157 (Fig. 2B). This result demonstrates that some of the effects of LL on leaf morphology do not require miR156/157.

**Figure 2.**
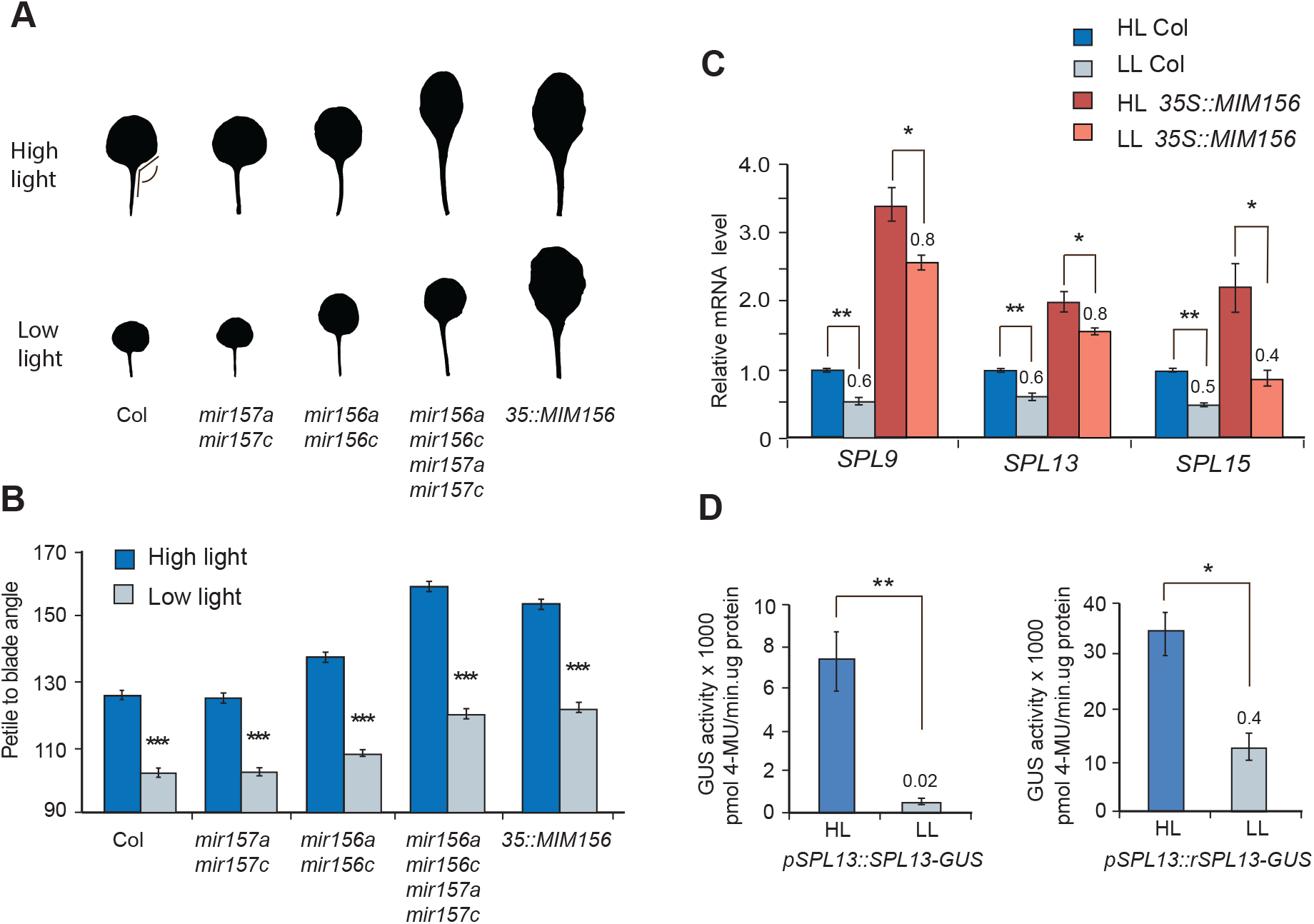
LL affects leaf morphology and SPL expression independently of miR156/157. (A) The morphology of leaf 1 under HL and LL in wild-type Col, in mutants lacking miR156 and/or miR157, and in plants constitutively expressing a miR156/157 target site mimic. (B) The petiole:blade angle of leaf 1&2 under HL and LL in wild-type Col and in genotypes deficient for miR156, miR157 and both miR156 and miR157. (C) The relative amount of SPL9, SPL13 and SPL15 transcripts in Col and *35S::MIM156* plants grown in HL and LL. Values are normalized to the transcript level in Col grown under HL. The ratio of the transcript abundance in LL relative to HL in each genotype is indicated. (D) MUG assays of GUS activity in transgenic plants expressing miR156-sensitive (left) and miR156-resistant (right) genomic constructs of SPL13-GUS under HL and LL. Note the difference in the scale of the Y axes. ***p<0.001, **p<0.01, *p<0.05, one-way ANOVA. RT-qPCR analyses and MUG assays are mean ± SEM from 3 biological replicates.

To determine if the effect of LL on leaf morphology might be mediated by a miR156/157-independent effect of LL on SPL gene expression, we measured the abundance of the SPL9, SPL13 and SPL15 transcripts in Col and *35S::MIM156* seedlings under HL and LL (Fig. 2C). As expected (He et al., 2018), under HL all three transcripts were elevated in *35S::MIM156* seedlings relative to Col. LL reduced the expression of all three transcripts in both Col and *35S::MIM156* plants, but appeared to have a slightly bigger relative effect on SPL9 and SPL13 in Col than in *35S::MIM156*, which is consistent with the elevated level of miR156 in LL Col plants.

miR156 represses SPL gene expression primarily at a translational level (He et al., 2018). Therefore, to obtain a more accurate picture of the relative importance of miR156-dependent vs. miR156-independent regulation of SPL activity, we examined the effect of light intensity on the expression of miR156-sensitive and miR156-resistant *pSPL13::SPL13-GUS* translational reporters. LL nearly completely repressed the expression of the miR156-sensitive *pSPL13::SPL13-GUS* reporter, and reduced the expression of the miR156-resistant *pSPL13::rSPL13-GUS* to about 0.4 of the level in HL plants (Fig. 2C and D). These results demonstrate that light intensity regulates *SPL* expression though both miR156-dependent and miR156-independent pathways.

### The response to light intensity is partially mediated by carbohydrate

Mutations in the chlorophyll oxygenase gene, *CHLORINA1* (*CH1*) (Oster et al., 2000), produce yellow-green plants that undergo delayed vegetative phase change (Yang et al., 2013), and thus resemble wild-type plants grown in LL. Wild-type plants grown in LL have about 60% of the amount of chlorophyll present in HL plants (Fig. 3A), whereas the *ch1-4* mutant has about 30% of the wild-type level chlorophyll (Fig. 3B). Previous work has suggested that *ch1-4* delays vegetative phase change because it reduces sugar production (Yang et al., 2013). To determine if the effect of LL on vegetative phase change is attributable to a reduction in the amount of sugar, we grew Col in HL and LL on media containing different concentrations of sucrose. 1% and 2% sucrose partially corrected the effect of LL on seedling growth, and 3% sucrose did not produce further changes (Fig. 3C). We then measured the abundance of miR156 and miR157 in Col grown in HL and LL in the absence or presence of 2% sucrose. Plants grown in LL had 1.8 times as much miR156 and 1.2 times as much miR157 as developmentally comparable plants grown in HL. 2% sucrose reduced the amount of miR156/miR157 in both HL and LL plants, but had a slightly greater effect on the level of these miRNAs in LL plants than in HL plants (Fig. 3D). We also examined the effect of 2% sucrose on the primary transcripts of *MIR156A* and *MIR156C* (Fig. 3E). Consistent with its effect on miR156, sucrose reduced the level of *pri-miR156a* and *pri-miR156c* more in LL plants than in HL plants (Fig. 3E). These results suggest that the effect of LL on the expression of miR156/miR157 is partially explained by the effect of LL on carbohydrate production.

**Figure 3.**
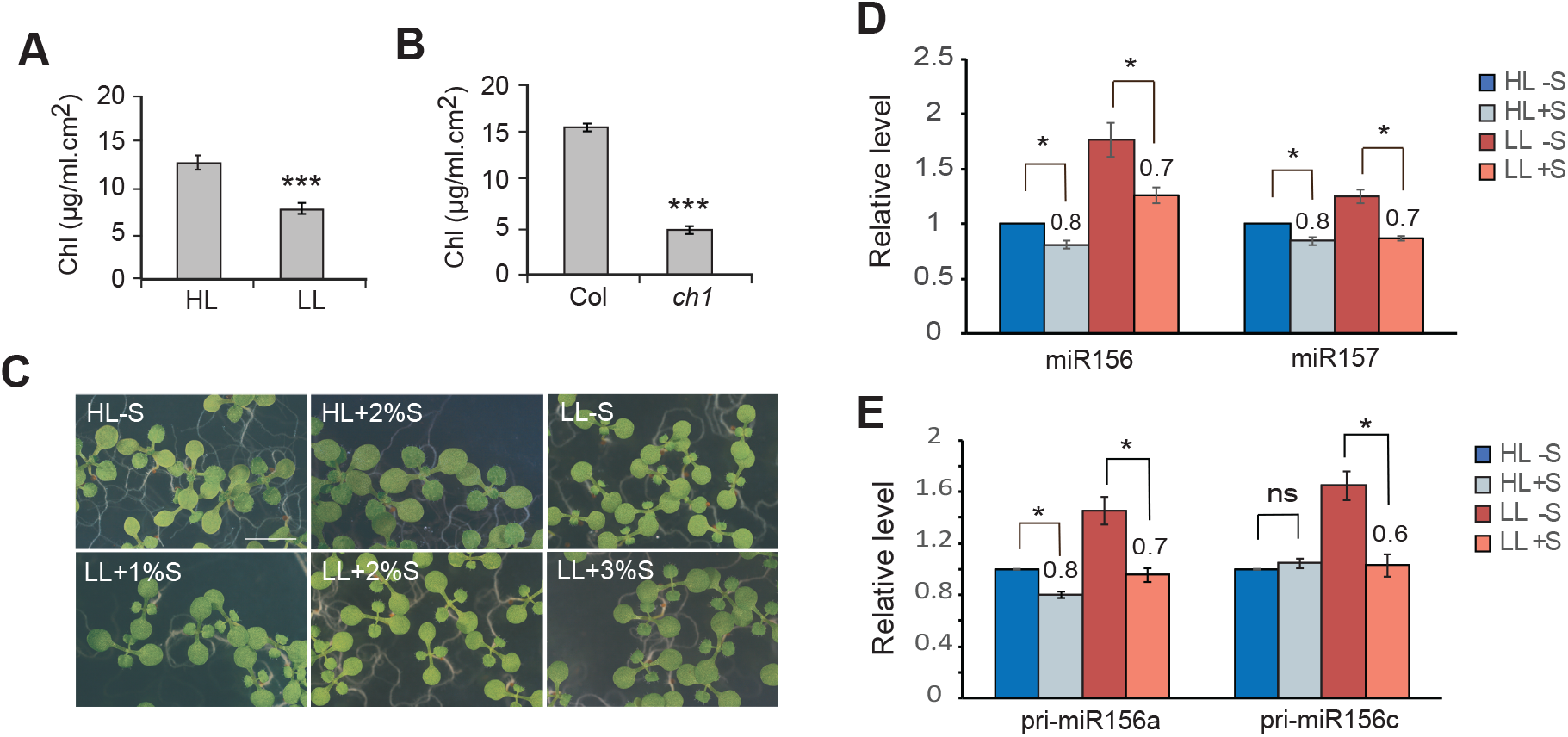
Sucrose partially corrects the de-repression of miR156 in LL. (A and B) Chlorophyll concentration in Col under HL and LL (A), and in Col and *ch1-4* under HL (B). Values are mean ± SEM from 4 biological replicates. (C) Col seedlings grown under HL and LL in the absence or presence of exogenous sucrose. (D) The effect of exogenous sucrose on the abundance of miR156 and miR157 in Col seedlings grown in LL. Values were normalized to the transcript leaves in Col under HL. The amount of transcript in LL relative to HL is shown. (E) The effect of exogenous sucrose on the abundance of pri-miR156a and pri-miR156c in Col seedlings grown in LL. Values were normalized to the transcript leaves in Col under HL. The amount of transcript in LL relative to HL is shown. ***p<0.001, *p<0.05, one-way ANOVA. Values are mean ± SEM from 3 biological replicates.

### Light intensity probably regulates miR156 expression independently of H3K27me3

Under HL, the down-regulation of *MIR156A* and *MIR156C* is mediated by the deposition of H3K27me3 at these loci (Pico et al., 2015; Xu et al., 2016). To determine if light intensity affects this process, we used chromatin immunoprecipitation (ChIP) to measure the level of H3K27me3 at *MIR156A*/*MIR156C* in HL and LL plants. The level of H3K27me3 at *MIR156A*/*MIR156C* was significantly lower in 2-week old LL plants than in 2-week old HL plants (Fig. 4A and B). However, in the case of *MIR156A*, this is probably explained by developmental delay caused by LL because 3-week old LL plants—which were at the same developmental stage as 2-week old HL plants—had nearly the same level of H3K27me3 at *MIR156A* as HL plants (Fig. 4A). In contrast, 3-week old LL plants had significantly less H3K27me3 at *MIR156C* than 2-week old HL plants (Fig. 4B). Thus, light intensity has a more significant effect on the deposition of H3K27me3 at *MIR156C* than at *MIR156A*. To determine if light intensity modulates the deposition of H3K27me3 by affecting carbohydrate production, we used ChIP to measure the level of H3K27me3 at *MIR156A*/*MIR156C* in two-week old plants growing in LL with or without sucrose. Sucrose significantly elevated the level of H3K27me3 at *MIR156A* (Fig. 4C) but had no effect on the level of H3K27me3 at *MIR156C* (Fig. 4D). In aggregate, these results indicate that LL and sugar can affect the level of H3K27me3 at *MIR156A* or *MIR156C* in ways that are consistent with the effect of these factors on the expression of these genes. However, the biological significance of these results is unclear because LL and sugar had different effects on the deposition of H3K27me3 at *MIR156A* and *MIR156C*, even though these factors affected the expression of these genes in the same way.

**Figure 4.**
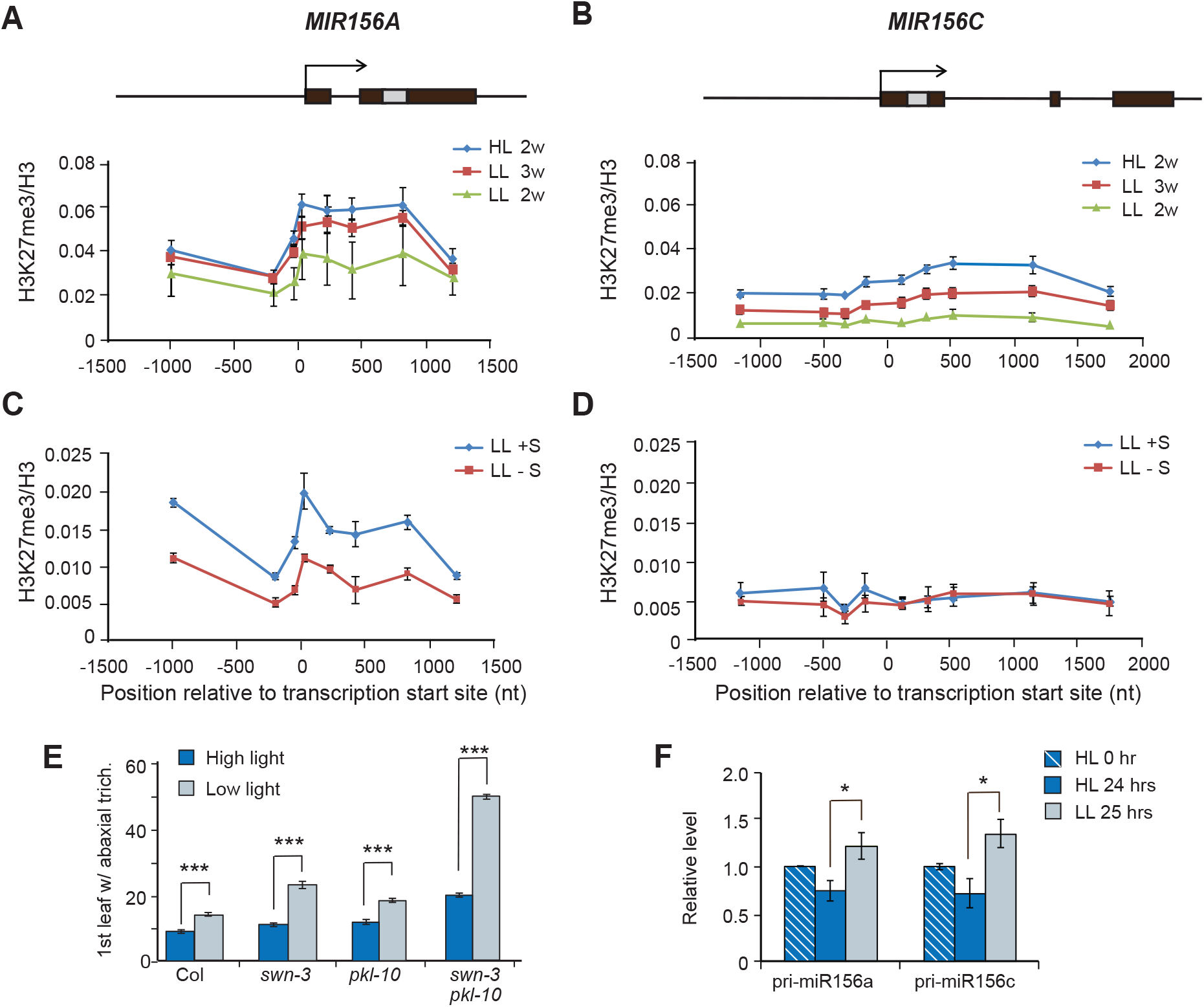
LL does not act by decreasing the deposition of H3K27me3 at *MIR156A* and *MIR156C*. (A and B) Schematic diagram of the genomic structure of *MIR156A* and *MIR156C* and ChIP-qPCR analysis of H3K27me3 at these loci in 2-week or 3-week old Col plants grown in HL or LL. Black boxes represent exons, the grey box represents the miR156 hairpin, and arrows indicate transcription start site. (C and D) ChIP-qPCR analysis of H3K27me3 at *MIR156A* (C) and *MIR156C* (D) in Col plants grown in LL with or without sucrose. (E) The first leaf with abaxial trichomes in Col, *swn-3, pkl-10*, and *swn-3 pkl-10* plants grown in HL and LL. n=20 to 24 for each genotype. (F) RT-qPCR analysis of the abundance of pri-miR156a and pri-miR156c transcripts in Col seedlings 24 hours after the transfer from HL to LL. ***p<0.001, *p<0.05, one-way ANOVA. Values in A-D and F are mean ± SEM from 3 biological replicates.

We next asked if the effect of LL on vegetative phase change requires H3K27me3. This repressive histone modification is deposited on target chromatin by the activity of Polycomb Repressive Complex 2 (PRC2) (Hennig and Derkacheva, 2009, 2009; Ito and Sun, 2009, 2009; Bouyer et al., 2011, 2011), one component of which is the histone methyl transferase, SWINGER (SWN) (Zhou et al., 2018; Shu et al., 2019). The CHD3-like chromatin remodeler, PICKLE (PKL), also promotes the deposition of H3K27me3, although the mechanism by which it does this is still unknown (Zhang et al., 2008; Zhang et al., 2012; Xu et al., 2016). Plants mutant for both *swn* and *pkl* have reduced levels of H3K27me3 at *MIR156A*/*MIR156C* and display elevated expression of both genes (Xu et al., 2016). To determine if changes in H3K27me3 are responsible for the effect of light intensity on vegetative phase change, we compared the phenotype of *swn-3 pkl-10* mutants under HL and LL. Previously we reported that under HL *swn-3, pkl-10*, and the *swn-3 pkl-10* double mutant undergo delayed vegetative phase change (Xu et al., 2016), and we were able to replicate this result (Fig. 4E). We also found that LL further delayed vegetative phase change in these mutants. This was particularly dramatic in the case of *swn-3 pkl-10*. In HL the appearance of abaxial trichomes was delayed by 5 leaves in *swn-3 pkl-10* (19.4 ± 0.3) compared to Col (7.8 ± 0.2); however, in LL, *swn-3 pkl-10* produced abaxial trichomes over 37 leaves later than Col (51.0 ± 0.9 vs. 13.1 ± 0.3) (Fig. 4E). This result demonstrates that a reduction in H3K27me3 makes plants hypersensitive to the effect of LL, but also suggests that effect of LL on vegetative phase change may not be mediated a LL-induced reduction in H3K27me3 because *swn-3 pkl-10* already has low levels of H3K27me3 at *MIR156A* and *MIR156C* (Xu et al., 2016).

H3K27me3 is usually deposited at a locus over several days or weeks (Jiang et al., 2008; Angel et al., 2011; Coustham et al., 2012; Yang et al., 2014; Xu et al., 2016). We reasoned that if LL de-represses *MIR156A*/*MIR156C* by reducing the deposition of H3K27me3, is it unlikely that a brief exposure to LL would cause a significant increase in the expression of these genes. To test this hypothesis, we grew Col in HL for 7 days and then compared the abundance of *pri-miR156a* and *pri-mirR156c* in the shoot apices of plants grown in HL for an additional 24 hours to plants transferred to LL for 24 hours. Plants grown in HL for an additional 24 hrs had less *pri-miR156a* and *pri-miR156c* than 7-day old HL plants, which is expected because the expression of these transcripts declines early in shoot development. In contrast, plants transferred to LL for 24 hrs had significantly more *pri-miR156a* and *pri-miR156c* than HL plants (Fig. 4F). Thus, LL has a relatively rapid effect on the transcription of *MIR156A*/*MIR156C*, suggesting that this effect is unlikely to be due to a decrease in H3K27me3 deposition.

### The response to low light intensity is not mediated by phytochrome or cryptochrome signaling

It has been reported that the SAS is regulated in part by a decrease in expression of miR156, mediated by exposure to FR light at the end of the day (Xie et al., 2017). We therefore decided to investigate if the R:FR and blue light signaling pathways contribute to the response of plants to LL. In Arabidopsis, the main photoreceptors for R and FR light are phyA and phyB, and the main blue light photoreceptors are CRY1 and CRY2 (Casal, 2013). To determine if these photoreceptors are required for the response to LL, we examined the phenotype of plants deficient for CRY1 and CRY2 (*hy4-B104 cry2-1*) and plants deficient for PHYA and PHYB (*phya-211 phyb-9*) (Fig. 5A). The vegetative morphology, as well as the timing of abaxial trichome production, in the *cry1 cry2* double mutant was essentially identical to WT under both HL and LL conditions (Fig. 5A). Consistent with previous reports (Reed et al., 1994; Devlin et al., 1996), under HL *phya phyb* double mutants were light green, and had an elongated hypocotyl, elongated internodes, small narrow leaves with elongated petioles, and flowered much earlier than WT. These double mutants also produced abaxial trichomes significantly earlier than WT plants under HL (Fig. 5A). *phya phyb* mutants had a similar if not identical morphology under LL, but produced abaxial trichomes significantly later under LL than under HL. However, the relative delay in abaxial trichome production in *phya phyb* mutants under LL vs HL was slightly less than in WT plants. These results indicate that CRY1 and CRY2 play no significant role in the response of plants to LL, and also suggest that PHYA and PHYB may little or no role in this response.

**Figure 5.**
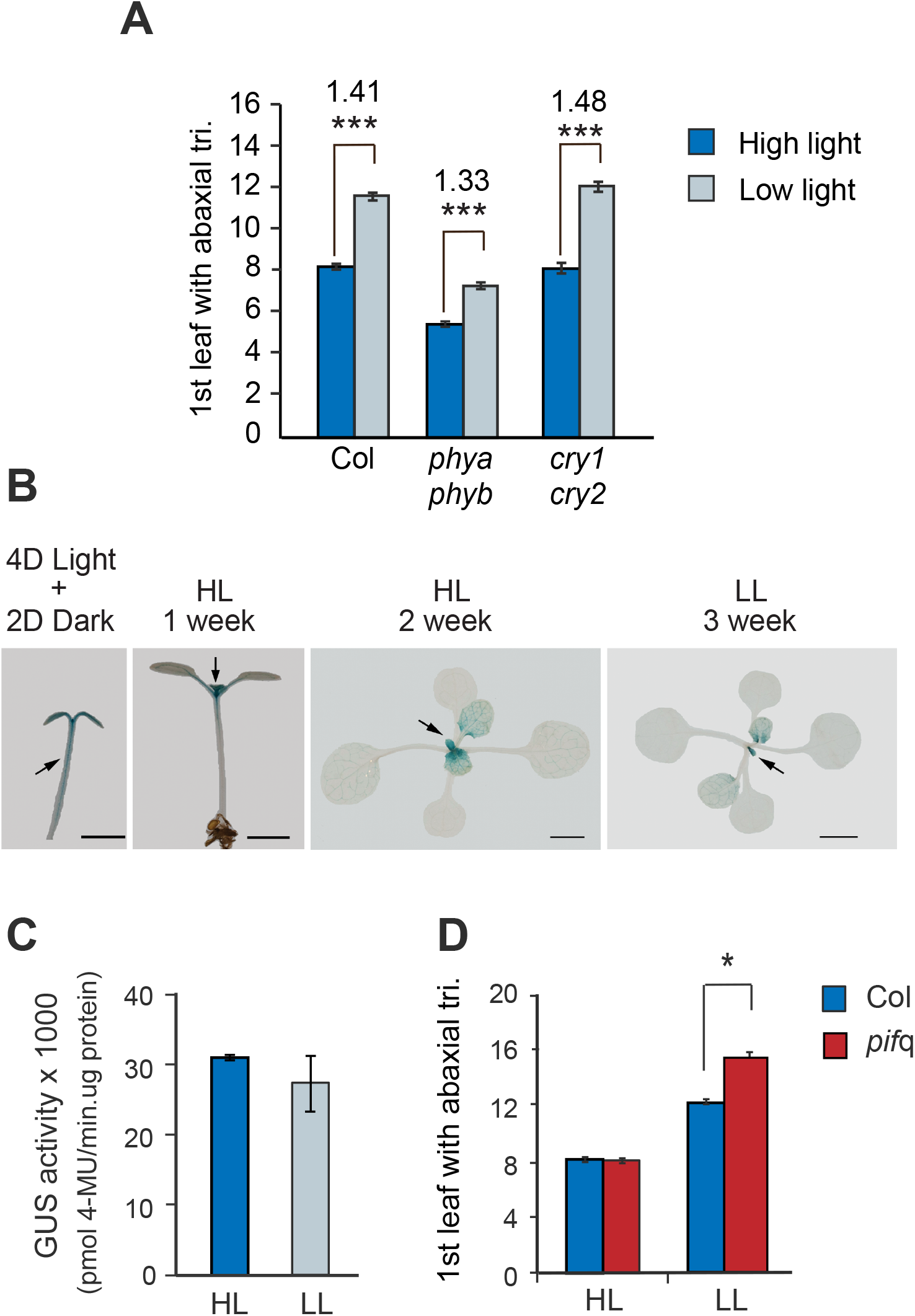
The red and blue light signaling pathways are not required for the effect of LL on vegetative phase change. (A) The effect of HL and LL on the first leaf with abaxial trichomes in Col and plants deficient for the red-light photoreceptors, phyA and phyB, and the blue light photoreceptors, CRY1 and CRY2. (B) Expression of PIF5::PIF5-GUS in seedlings and rosette leaves in HL and LL. (C) Quantitative MUG assay of PIF5::PIF5-GUS activity in the HL and LL rosettes shown in (B). (D) First leaf with abaxial trichome production in Col and *pif*q. ***p<0.001, *p<0.05, one-way ANOVA.

To further explore the role of blue and red light in the STS, we examined the effect of LL on genes that mediate the effect of these stimuli. Red and blue light control plant growth by regulating the expression of a family of basic helix-loop-helix (bHLH) transcription factors known as PHYTOCHROME INTERACTING FACTORS (PIF) (Pedmale et al., 2016). PIF proteins accumulate in the presence of a low R:FR ratio and low intensity blue light, and promote the expression of genes that generate the SAS. Plants mutant for *pif4* and *pif5*, as well as the *pif1 pif3 pif4 pif5* quadruple mutant (*pif*q), do not display the SAS under low R:FR light or low intensity blue light, and resemble plants grown in HL (Leivar et al., 2012). If PIF proteins promote the STS as well as the SAS, the expectation is that that these proteins will be over-expressed under low light intensity and that loss of PIF gene expression will accelerate the onset of the adult phase.

To test these hypotheses, we first examined the effect of light intensity on the expression of *PIF5* in seedlings and rosette leaves. Consistent with previous observations (Pedmale et al., 2016), the *PIF5::GUS* transcriptional reporter was highly expressed in the hypocotyl, shoot apex and cotyledons of seedlings germinated in HL for 4 days and transferred to the dark for 2 days, but was only expressed in the shoot apex of seedlings grown continuously in HL (Fig. 5B); thus, *PIF5::GUS* expression in the hypocotyl is repressed by HL. In 2-week old plants grown continuously in HL, *PIF5::GUS* was strongly expressed in young leaf primordia but was restricted to the vascular tissue of fully expanded leaves. It had a similar expression pattern in 3-week old plants grown in LL, which were developmentally comparable to the 2-week old HL plants (Fig. 5B). MUG assays revealed no significant difference in the amount of GUS activity in the shoot apex of plants grown in HL and LL (Fig. 5*C*). Therefore--in contrast to its expression in the hypocotyl--PIF5 expression in rosette leaves is insensitive to light intensity. We next examined the effect of the *pif*q mutant on abaxial trichome production under HL and LL conditions. There was no significant difference in the timing of abaxial trichome production in wild type and *pif*q mutant in HL; however abaxial trichome production was delayed more significantly in LL in the *pif*q mutant than in Col (Fig. 5D). This latter result implies that PIF proteins repress the response of plants to LL, which is the opposite of their function in the response to low R:FR light (i.e the SAS), which they promote. Together, these results indicate that LL delays vegetative phase change by a mechanism that is distinct from the mechanism involved in the SAS.

## Discussion

The STS is typically considered to consist of a set of traits that enable a plant to fix carbon above the photosynthetic compensation point under low light intensity (5); in other words, it is viewed as a physiological adaptation to low light intensity. This begs the question of how LL promotes the expression of these traits, and why the STS includes traits that have no obvious connection to photosynthesis. Our results suggest that the answer to these questions lies in the effect of LL on the expression of genes that regulate vegetative phase change, specifically, miR156/miR157, and their targets, the SPL family of transcription factors. This conclusion is supported by the observation that juvenile leaves of Arabidopsis, maize, and poplar are photosynthetically more efficient under LL than adult leaves (Lawrence et al., 2020) and by the results of this study.

We found that LL prolongs the duration of the juvenile phase and that it does so by both increasing the abundance of miR156 and by decreasing the transcription of the SPL genes regulated by miR156. These effects appear to be mediated by a reduction in photosynthesis, as well as by other unknown factors. Importantly, the effect of LL on vegetative phase change does not require the photoreceptors that mediate the response of Arabidopsis to red and blue light, specifically, phyA, phyB, CRY1 and CRY2, or the PIF transcriptions factors that mediate the effects of these photoreceptors. This supports the conclusion that LL regulates vegetative phase change by a different mechanism, or set of mechanisms, than the mechanism responsible for the SAS. Further supporting this conclusion, the traits associated with the SAS are opposite to the traits associated with the STS. Specifically, in Arabidopsis, exposure to a low R:FR ratio or end-of-day FR light produces elongated internodes, and light green elongated leaves with long petioles, and accelerated abaxial trichome production, whereas LL produces normal length internodes, and rounded, dark green leaves with normal length petioles and delayed abaxial trichome production.

Evidence that LL operates through both miR156-dependent and miR156-independent mechanisms is provided by the effect of LL on plants deficient for miR156. Specifically, we found that miR156 is essential for the effect of LL on abaxial trichome production, but is not essential for its effect on leaf shape. We also found that the expression the direct targets of miR156 is reduced by LL even in a genetic background with little or no functional miR156 and miR157. Finally, we found that LL reduces the expression of a miR156-insensitive translational reporter for SPL13. Together, these observations suggest that specific phenotype of plants grown in LL may depend on the SPL genes that regulate particular traits and on the relative importance of post-transcriptional (i.e. miR156-dependent) vs. transcriptional regulation for the expression of these SPL genes. This scenario could explain why the phenotype of plants growing under low light intensity is similar but not necessarily identical to the phenotype of juvenile shoots growing under high light intensity (Jones, 1995).

The observation that exogenous sucrose partially corrects the effect of LL on the expression of *MIR156A* is consistent with previous studies demonstrating that sugar accelerates vegetative phase change (Yang et al., 2013; Yu et al., 2013), and supports the hypothesis that LL delays vegetative phase change through its effect on photosynthesis. However, this is not the only explanation for the effect of LL on vegetative phase change because sugar did not correct the LL-induced increase in the expression of *MIR156C* or the LL-induced reduction in H3K27me3 at this gene. Although this decrease in H3K27me3 may contribute to the increase in *MIR156C* expression, it is unlikely to be the only factor involved in this response because *MIR156C* expression increases relatively quickly in response to LL. Together, these results suggest that LL regulates the expression of miR156 by reducing photosynthesis, and by several light-sensitive processes or pathways that operate independently of photosynthesis.

The observation that shade intolerant species are able to display features characteristic of the STS when grown in LL suggests that the genes responsible for the STS are not unique to shade tolerant species. Interspecific comparisons of plants with strongly differentiated juvenile and adult phases have suggested that the morphology of juvenile shoots is an adaptation to LL environments (Day, 1998), and this is supported by studies in species with less highly differentiated juvenile and adult phases (Lusk, 2004; Lusk et al., 2008). Additional support for this conclusion comes from the observation that the photosynthetic properties of juvenile leaves in English ivy (*Hedera helix*) resemble those of leaves adapted for growth in LL (Bauer and Bauer, 1980), and from a recent study demonstrating that juvenile leaves, as well as leaves over-expressing miR156, have a significantly higher maximum rate of photosynthesis under LL than adult leaves and that the rate of photosynthesis in these leaves is less sensitive to LL than the rate of photosynthesis in adult leaves (Lawrence et al., 2020). These studies, and the results presented here, suggest that the juvenile phase is naturally adapted for LL conditions, and further suggest that the STS is regulated by the vegetative phase change pathway.

## Materials and Methods

### Plant Material and Growth Conditions

All mutants used in this study were in a Columbia background. *mir156a-2, mir156c-1, mir157a-1, mir157c-1, 35S:MIM156, chlorina1-4, swn-3, pkl-10*, and *swn3 pkl-10* have been described previously (Wang et al., 2009; Yang et al., 2013; Xu et al., 2016; He et al., 2018). Seeds of the *phya-211 phyb-9* and *cry1(hy4-B104) cry2-1* double mutants were provided by Dr. Meng Chen (U. of California, Riverside). The *phyb-9* lines lack the mutation in the *VENOSA-4* gene that is present in the original *phyb-9* stock. The *pif1-1 pif3-7 pif4-2 pif5-3* (*pifq*) quadruple mutant was obtained from the Arabidopsis Resource Center and the *PIF5::GUS* line was provided by Dr. Joanne Chory (Salk Institute, La Jolla, CA). Plants were grown in Conviron growth chambers in short days (10 hrs light:14 hrs dark) at 22°C, with illumination provided by a combination of Sylvania warm white (4000 K) and Interlectric WS Gro-lite T8 fluorescent bulbs. High light conditions consisted of an overall intensity of 180 µmol m^-2^ s^-1^ [15. 7 µmol m^-2^ s^-1^ blue (450-550 nm), 59 µmol m^-2^ s^-1^ red (610-740 nm), 9.4 µmol m^-2^ s^-1^ far red (710-850 nm)]. Low light conditions consisted of an overall intensity of 62 µmol m^-2^ s^-1^ [5 µmol m^-2^ s^-1^ blue (450-550 nm), 20 µmol m^-2^ s^-1^ red (610-740 nm), 3.75 µmol m^-2^ s^-1^ far red (710-850 nm)]. For sugar treatment, plants were grown in 1/2 MS plates with or without 1%, 2%, 3% sucrose as indicated.

### Quantitative RT-PCR

RNA for RT-qPCR analysis was isolated from shoot apices containing leaves less than 1 mm in length. HL or LL-grown plants were compared at the same developmental stage. For example, the SL1+ apices were harvested by removing the two cotyledons when leaf primordia 1 and 2 had just emerged. SL3+ apices were harvested when leaf primordium 3 was just visible, and the cotyledons and leaf 1 and 2 were removed. Total RNA was isolated using TRIzol method, followed by DNase treatment (Ambion) according to the manufacturer’s instructions. The primers used for this study were the same as the primers in Xu et al. (2016).

### Chlorophyll Concentration

Leaf punches were collected from the middle of lamina (leaf 3 or leaf 4) and ground in a mortar and pestle. The ground tissue was suspended in 80% acetone and centrifuged briefly. The optical density of the supernatant was measured at 645 nm and 663 nm using a spectrophotometer, and chlorophyll concentration was calculated as (OD_645_ x 20.2 + OD_663_ x 8)/ml·cm^2^).

### Chromatin Immunoprecipitation

Chromatin was isolated from Col plants at the same developmental stage and immunoprecipitated with antibodies to H3 or H3K27me3 as previously described (Xu et al., 2016). DNA sequences were analyzed by qPCR, using the primers for *MIR156A* and *MIR156C* described in (Xu et al., 2016).

## ACKNOWLEDGEMENTS

We gratefully acknowledge helpful discussions with Dr. Meng Chen (U. of California, Riverside) and members of the Poethig laboratory.

## Figure Legends

**Figure S1.** LL delays vegetative phase change. (A) RT-qPCR analysis of SPL9, SPL13, and SPL15 transcripts in L5+ shoot apices from plants grown in HL and LL. *p<0.05, one-way ANOVA. Values are mean ± SEM from 3 biological replicates. (B) The morphology and pattern of abaxial trichome production in Col, *mir156a mir156c, mir157a mir157c, mir156a mir156c mir157a mir157c*, and *35S::MIM156* plants grown in HL and LL conditions. Grey = no abaxial trichomes. Black = abaxial trichomes.

